# Multi-objective Bayesian Optimization with Heuristic Objectives for Biomedical and Molecular Data Analysis Workflows

**DOI:** 10.1101/2022.06.08.495370

**Authors:** Alina Selega, Kieran R. Campbell

## Abstract

Many practical applications require optimization of multiple, computationally expensive, and possibly competing objectives that are well-suited for multi-objective Bayesian optimization (MOBO) procedures. However, for many types of biomedical data, measures of data analysis workflow success are often heuristic and therefore it is not known *a priori* which objectives are useful. Thus, MOBO methods that return the full Pareto front may be suboptimal in these cases. Here we propose a novel MOBO method that adaptively updates the scalarization function using properties of the posterior of a multi-output Gaussian process surrogate function. This approach selects useful objectives based on a flexible set of desirable criteria, allowing the functional form of each objective to guide optimization. We demonstrate the qualitative behaviour of our method on toy data and perform proof-of-concept analyses of single-cell RNA sequencing and highly multiplexed imaging datasets.

## Introduction

The analysis of high-dimensional biological data is often exploratory and unsupervised. For example, gene expression data may be subject to clustering algorithms to find groups representative of meaningful biological variation. For assays that profile at the patient level, these clusters may represent novel disease subtypes, while for assays at the single-cell level, they may represent novel cell types.

Despite the importance of these methods, there is no “one-size-fits-all” approach to the analysis of such data. Instead, there is a myriad of different possible parameter combinations that govern these workflows and lead to variations in the results and interpretation. For example, in the analysis of single-cell RNA-sequencing (scRNA-seq) – a technology that quantifies the expression profile of all genes at singlecell resolution – a common analysis strategy is to cluster the cells to identify groups with biological significance. However, each workflow for doing so has variations with respect to data normalization, cell filtering strategies, and the choice of clustering algorithm and parameters thereof. Changes to these algorithm and parameter choices produce dramatically different results (1, 2) and there is no ground truth available. This motivates an important question: how do we optimize these workflows such that the resulting exploratory analysis best reflects the underlying biology?

In the adjacent field of supervised machine learning (ML), such optimization over workflows has largely been tackled from the perspective of automated ML (AutoML, (3)). This comprises a diverse set of methods such as Bayesian optimization (4) and Neural Architecture Search (5) that attempt to optimize the success of the model with respect to one or more hyperparameter settings. In this context, success is defined as the model accuracy on a held out test set, though can also correspond to the marginal likelihood of the data given the model and hyperparameters.

However, in the context of exploratory analysis of genomic data, existing AutoML approaches face three challenges. Firstly, they are almost exclusively unsupervised, meaning there is no notion of accuracy on a test set we may optimize with respect to. Secondly, the majority of methods are not generative probabilistic models (6) so it is impossible to optimize with respect to the marginal or test likelihood. Finally, the objectives used to optimize a workflow are numerous, conflicting, and can be highly subjective, due to often being heuristics.

This is demonstrated by attempts to benchmark clustering workflows of scRNA-seq data. As said above, there are many parameters that must be set, e.g. which subset of genes and clustering algorithm to use, along with such parameters as resolution in the case of community detection (1). However, there is no quantitative way to choose which parameter setting is “best” and so the community turns to a number of heuristic objectives to quantify the performance of a workflow. For example, Cui et al. (7) attempt to optimize the adjusted Rand index (ARI) with respect to expert annotations and a heuristic based around downsampling rare cell types while minimizing runtime. Germain et al. (1) similarly consider the ARI but also the average silhouette width to maximize cluster purity. Zhang et al. (8) consider a range of heuristics including agreement with simulated data and robustness to model misspecification.

However, given that these objectives are all heuristic and open to user preference, there is no guarantee that all of them are *useful* and have maxima that align with the meta-objective at hand, which in the above example is the ability to identify a biologically relevant population of cells. Conversely, some heuristic objectives may be *non-useful* – they are largely noisy and attribute nothing to the overall optimization problem by not aligning with a meta-objective. This motivates the central question we attempt to address: how can we adapt AutoML approaches to optimize unsupervised workflows over multiple heuristic objectives that are frequently subjective and conflicting?

To begin to tackle this question, we introduce MANATEE (Multi-objective bAyesiaN optimizAtion wiTh hEuristic objEctives). The key idea is that by considering a linear scalarization as a probabilistic weighting over (heuristic) objective inclusion, we may up- or downweight an objective based on desirable or non-desirable behaviours of its posterior functional form. Consequently, rather than returning the full Pareto front that may include points (parameter combinations) that maximize potentially non-useful heuristic objectives, we automatically concentrate on a useful region. The main contributions presented here are:

1. Introduce the concept of *behaviours* 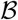 of the posterior functional form of the surrogate objective function **f** that are desirable if a function is useful for overall optimization.
2. Suggest a set of such behaviours that may be inferred from the posterior of a multi-output Gaussian process, if used as the surrogate function.
3. Build upon previous MOBO procedures to compute the distribution of scalarization weights 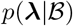 with resulting optimizations returning Pareto optimal points.
4. Devise a set of experiments based on the analysis of real molecular imaging and transcriptomic data and show that the proposed procedure compares favourably to existing approaches.

## Background

### Bayesian optimization

Bayesian optimization (BO, see (9) and references therein) attempts to optimize a function 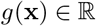 for some 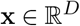 that is, in some sense, expensive to evaluate and for which derivative information is not available, precluding gradient-based approaches. Applications of BO have become popular in the tuning of ML hyperparameters (10) and indeed entire workflows (11) due to the expensive nature of re-training the models.

At their core, BO approaches propose a surrogate function *f* defined on the same range and domain as *g* that may be searched efficiently to find points *x* that either maximize *g*, reduce uncertainty about *f*, or both. This leads to the concept of an acquisition function 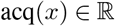 that may be optimized to find the next *x* at which *g* may be evaluated. While multiple acquisition functions have been proposed, including probability of improvement and expected improvement, here we focus on the Upper Confidence Bound (UCB) defined as

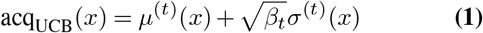

where *μ*^(*t*)^(*x*) and *σ*^(*t*)^(*x*) are the posterior mean and standard deviation of *f* at *x* after *t* acquisitions from *g*, while *β_t_* is a hyperparameter that controls the balance between exploration and exploitation. While there are many possible choices for the surrogate function *f*, including deep neural networks (12), a popular choice is a Gaussian process due to its principled handling of uncertainty and capacity to approximate a wide range of functions.

### Gaussian processes

Gaussian processes (GPs) (13) define a framework for performing inference over nonparametric functions. Let *m*(*x*) be a mean function and *k*(*x, x*′) a positive-definite covariance function for 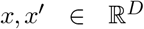. We define *f*(*x*) to be a Gaussian process denoted 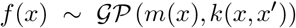 if for any finitedimensional subset 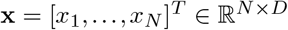, the corresponding function outputs **f** = [*f*(*x*_1_), …, *f*(*x_N_*)] follow a multivariate Gaussian distribution 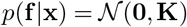, where **K** is the covariance matrix with entries (**K**)*_ij_* = *k*(*x_i_, x_j_*) and we have assumed a zero-mean function without loss of generality. The kernel fully specifies the prior over functions, with one popular choice we use throughout the paper being the *exponentiated quadratic* kernel 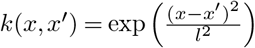. It is common to model noisy observations **y** via the likelihood *p*(**y**|**f**), which when taken to be 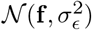 leads to the exact marginalization of **f**.

### Multi-output Gaussian processes

Gaussian processes may be extended to model *K* distinct outputs^1^ via the functions 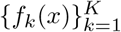. One construction is to model the full covariance matrix as the Kronecker product between the *K* × *K* inter-objective covariance matrix **K**^IO^ and the data covariance matrix:

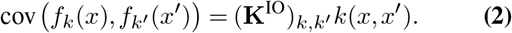

Here the kernel hyperparameter *l* is shared across objectives, though in the following we model objective-specific observation noises *ϵ_k_*.

### Multi-objective optimization

Multi-objective optimization attempts to simultaneously optimize *K* objectives *g*_1_(*x*), …, *g_K_*(*x*) over 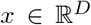, which is common in many real-world settings. However, it is rare in practice to be able to optimize all *K* functions simultaneously and instead is common to attempt to recover the *Pareto front*. We say a point *x*_1_ is *Pareto dominated* by *x*_2_ iff *g_k_* (*x*_1_) ≤ *g_k_*(*x*_2_) ∀*k* = 1, …, *K* and ∃*k* ∈ 1, …, *K* s.t. *g_k_*(*x*_1_) < *g_k_*(*x*_2_). A point is said to be *Pareto optimal* if it is not dominated by any other point. The *Pareto front* is then defined as the set of Pareto optimal points, which intuitively corresponds to the set of equivalently optimal points given no prior preference between objectives.

### Scalarization functions

One popular approach to multiobjective optimization is the use of *scalarization functions* (see (14) for an overview). A scalarization function *s*_**λ**_(**g**(*x*)) parameterized by **λ** takes the set of *K* functions **g**(*x*) = [*g*_1_(*x*), …, *g_K_*(*x*)] and outputs a single scalar value to be optimized in lieu of **g**(*x*). It can be shown (15) that if *s*_**λ**_ is monotonically increasing in all *g_k_*(*x*) then the resulting optimum *x*\* lies on the Pareto front of **g**.

While many scalarization functions exist, one popular choice is the linear scalarization function 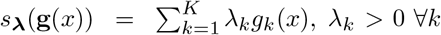. This has the intuitive interpretation that each λ*_k_* corresponds to the weight assigned to function *k*, with a larger relative value pulling the optimum of *s*_**λ**_ towards the optimum of *g_k_*.

### Hypervolume improvement

Another multi-objective optimization approach relies on the notion of *hypervolume* (HV), the volume of the space dominated by an approximate Pareto front and bounded from below by a reference point, which current work assumes to be known by the practitioner (16). HV is used as a metric to assess the quality of a Pareto front and is sought to be maximized in the optimization. HV can be efficiently computed by box decomposition methods (17), allowing one to compute the HV improvement (HVI) for a new set of points (18).

### Multi-objective Bayesian optimization

MOBO procedures tackle the scalarization-based multi-objective optimization as above but under the same conditions as BO, where each evaluation of *g_k_*(*x*) is expensive and derivative information is unavailable. An example method is ParEGO (19), which randomly scalarizes objectives with augmented Chebyshev scalarization and uses expected improvement. It was recently extended to *q*NParEGO (20), which supports parallel and constrained optimization in a noisy setting. Unlike hypervolume-based methods which can struggle with > 5 objectives (21), *q*NParEGO is more suited for such problems.

Paria et al. (15) propose a MOBO procedure that, rather than maximizing *s*_**λ**_ for a single **λ**, constructs a distribution *p*(**λ**) and minimizes the expected pointwise regret,

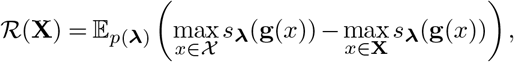

where 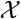 is the feature space of *x* and **X** is the subset of 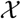 lying on the Pareto front to be computed. The exact region of the Pareto front to be considered is governed by *p*(**λ**) and the authors provide a bounding box procedure for the user to select *p*(**λ**), akin in the case of a linear scalarization to asserting *a priori* which objectives *k* are important. However, to our knowledge, no MOBO approach has proposed a *p*(**λ**|·), inferred from either the data or the posterior over functions, that adaptively up- or downweights objectives based on desirable properties.

For hypervolume-based methods in the MOBO setting, the posterior distribution of the surrogate model can be used to compute the expected hypervolume improvement (EHVI) (20). As EHVI assumes noise-free observations and can be affected in the presence of noise, recent work introduced noisy hypervolume improvement (NEHVI), which uses the expectation of EHVI under the posterior distribution of the surrogate function values given noisy observations (16). NEHVI is more robust to noise than other hypervolume-based MOBO methods, is equivalent to EHVI in the noiseless setting, and its parallel formulation (*q*NEHVI) achieves computational gains and state-of-the-art performance in large batch optimization (16).

### Applications of AutoML in genomics

Despite the prevalence of ML applications in genomics, AutoML methods have received surprisingly little attention. The GenoML project (22) provides a Python framework centered on open science principles to perform end-to-end AutoML procedures for supervised learning problems in genomics. Auto-GeneS (23) develops a multi-objective optimization framework for the selection of genes for the deconvolution of bulk RNA-sequencing. However, to our knowledge, there is no work that tackles the general problem of optimizing bioinformatics and genomics workflows in the absence of well-defined objective functions. In contrast, there are multiple BO techniques that allow a user to express a preference between solutions (24). While these could have exciting applications in genomics, we assume such information is not available here.

## Multi-objective Bayesian optimization over heuristic objectives

### Setup

We assume we have access to *K* noisy, heuristic objectives that at acquisition step *t* return a measurement *y_kt_* for an input location 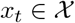, where 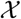 is a compact subset of 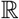 on [*a, b*]. We introduce surrogate functions *f_k_*(*x*) that we model with a multi-output GP as described above with a full kernel given by Equation 2. We use a linear scalarization function over objectives 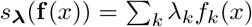 and ultimately seek to maximize 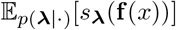. The next point to query *x*_*t*+1_ is chosen by maximizing the expectation of the acquisition function. For this we propose two approaches: (i) the expectation of the scalarization of the single-objective acquisition function of each objective 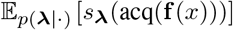 as per (15) and (ii) the expectation of the single-objective acquisition function of the scalarized objectives 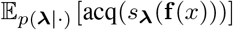 (derived in Supplementary Note 3). We denote these as SA (scalarized acquisition) and AS (acquisition of scalarized), respectively, and use the UCB single-objective acquisition function as per Equation 1. We wish to set *p*(**λ**|·) to upweight objectives that are inferred as useful based on desirable properties learned from the data.

### Desirable heuristic objective behaviours

We begin by considering what properties of a given heuristic objective *f_k_*(*x*) may be considered desirable that would lead to upweighting of that objective in our overall optimization procedure. While many are possible, we suggest three (Figure 1):

1. **Explainability:** *f_k_*(*x*) covaries significantly with *x* (i.e. is explained by *x*). The justification here is that the practitioner has selected heuristic *k* assuming it will provide insight into the choice of *x*, so if there is no correlation then it should be downweighted. Given that the data have been scaled to empirical variance 1, 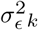 represents the proportion of variance unexplained by *f_k_* so we define 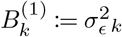.
2. **Inter-objective agreement:** *f_k_* shares a similar functional form with 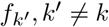, with the intuition that it is useful for practitioners to find regions of the input space where multiple heuristics agree. After fitting the multi-output GP, (**K**^IO^)_*k, k′*_ defines the covariance between objective *k* and *k*′ for *k* ≠ *k*′ and (**K**^IO^)_*k, k*_ defines the variance of objective *k*. We therefore introduce the inter-objective agreement behaviour as

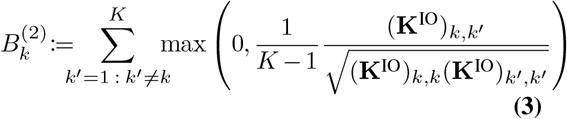 The intuition is that 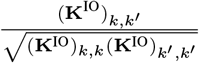 represents the correlation between objectives *k* and *k*′ so 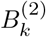 represents the average correlation with other objectives while not penalizing negative correlation worse than no correlation.
3. **Maximum not at boundary:** Within 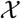, *f_k_* has a maximum that is not at the boundary of *x*. The justification is that if the maximum is at the boundary, *f* may be unbounded by increasing |*x*| and/or there is a mismatch between the practitioner’s expectations of the domain of 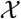 and the behaviour of *f*. Since the derivative of a GP is also a GP, we may identify whether a stationary point exists in 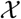 by searching for the zeros of the posterior mean derivative 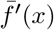. We therefore define 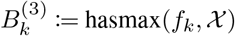, where hasmax returns 1 if *f_k_* has a maximum on 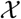 and 0 otherwise by evaluating the derivatives of the posterior mean of the multi-output GP (derived in Supplementary Note 4).

**Fig. 1.**
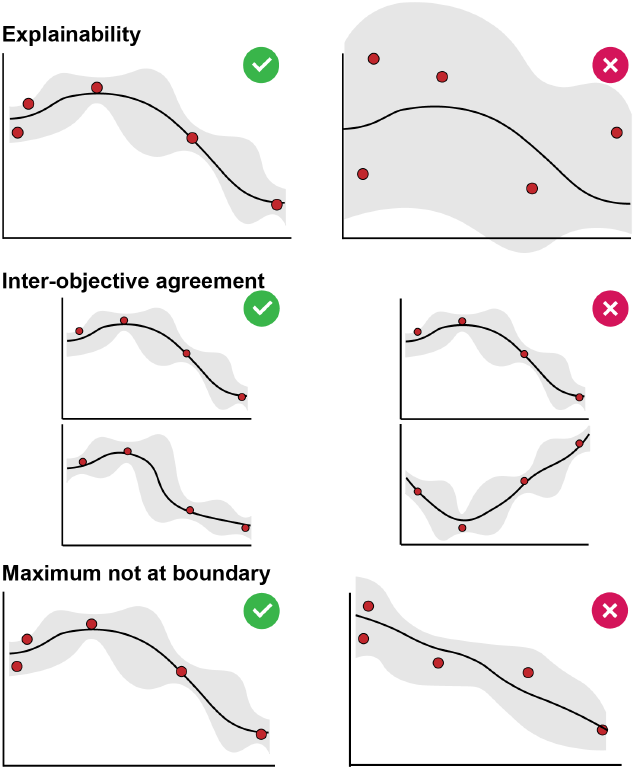
Examples of desirable behaviours.

### Incorporating desirable behaviours into scalarization weights

We next consider how to use the set of behaviours 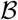 to parameterize the scalarization probabilities 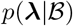. We assume that *λ_k_* is a binary variable ∀*k* that corresponds to whether objective *k* is *useful* or otherwise, with *p*(λ*_k_* |**B**_*k*_) given by a Bernoulli distribution. While this construction initially appears restrictive, it has two desirable properties that maintain its generality (proofs presented in Supplementary Note 5):

#### Theorem 1

If 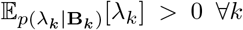, the solution to 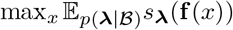 lies on the Pareto front of **f**.

#### Theorem 2

For some 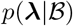, any point *x*\* on the Pareto front of **f** is reachable as a maximizer of 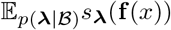.

However, how to construct 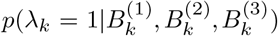 directly is non-obvious. Instead, we ask how would *each* objective behaviour appear if we knew that objective was useful or otherwise? These allow us to specify 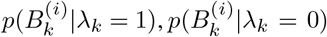 for *i* = 1, 2, 3 and combine with a prior *p*(λ*_k_* = 1) = 1 – *p*(λ_*k*_ = 0) to compute 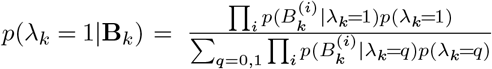. With these considerations, we suggest distributions for 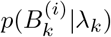; however, we emphasize that these are suggestions only and there are many possible that would fit the problem.

#### Explainability

For 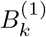, the explainability of objective *k* (i.e. the proportion of variance unexplained by that function), we assume that if that objective is desirable (λ*_k_* = 1) then the lower the observation noise, the better and in the non-desirable case (*λ_k_* = 0), higher noise is expected. Given the lack of additional assumptions, we appeal to the principle of parsimony and propose a linear relationship of the form:

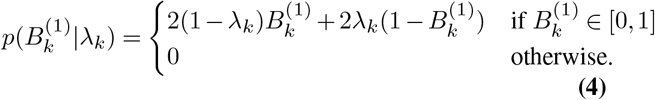

#### Inter-objective agreement

For inter-objective agreement 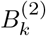, we propose reversed likelihoods to Equation 4 given the reasoning that high inter-objective correlation should be more likely under a desirable objective and vice-versa for a non-desirable one, and again a linear relationship is the most parsimonious.

#### Maximum not at boundary

We propose 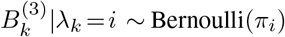 where π_0_, π_1_ are user-settable hyper-parameters. This means that conditioned on an objective being useful (or otherwise), there is a fixed probability of that objective containing a maximum in the region.

### MANATEE

Putting these steps together results in the MANATEE framework, an iterative MOBO procedure as outlined in Figure 2. First, the objectives are evaluated at a set of input locations randomly chosen on the parameter space. Second, the multi-output GP surrogate function with covariance given by Equation 2 is fitted to all objectives. Then, the objective behaviours 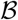 are computed from the surrogate function and the distributions over objective weights are updated. Finally, the updated acquisition function is optimized, guiding acquisition of the next point. The procedure is repeated for a predetermined number of steps. The overall “best” point to be used for downstream analysis may be chosen as that which maximizes the scalarized surrogate function.

**Fig. 2.**
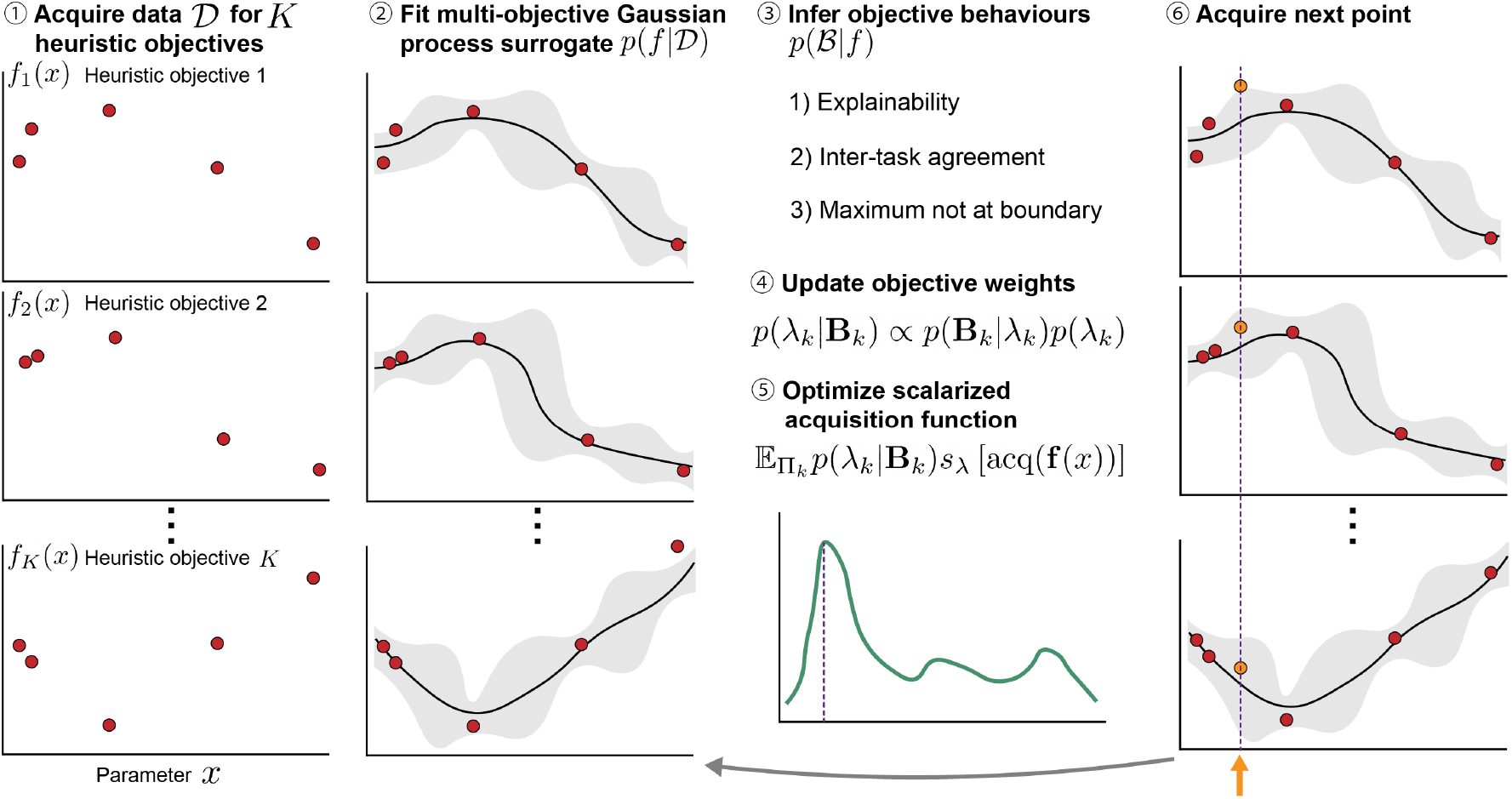
Multi-objective Bayesian optimization with heuristic objectives.

### Baselines for experiments

We contrast our method against two baselines and two existing approaches designed for MOBO of noisy objectives: (i) *Random acquisition*: draw *x_t_* ~ Unif(0, 1) at each iteration, (ii) *Random scalarization*: use identical surrogate and acquisition functions as MANATEE to sample *x_t_* but draw *λ_k_* ~ Unif(0, 1) rather than conditional on 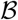, (iii) *q*NEHVI (16) with approximate hypervolume computation to facilitate inference over > 5 objectives, and (iv) *q*NParEGO (20).

When a meta-objective *h*(*x_t_*) is available at every iteration *t* = 1, …, *T* with overall maximum 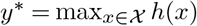, to compare among approaches we compute the following metrics: (i) *Cumulative regret*: 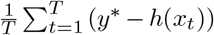, (ii) *Full regret*: 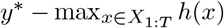, and (iii) *Bayes regret*: 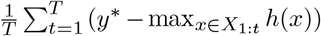, where *X*_1:*t*_ is the set of *x* acquired up to time *t*. Of these, we place most emphasis on cumulative regret as it quantifies how close each method gets to the optimal solution on average. In contrast, the full and Bayes regret quantify how close the “best” acquired point gets to *y*\* as measured by the max over *h* of all points acquired so far; however, since the meta-objective *h* is in general inaccessible for our problem setup (and only used for method comparison), it is impossible to quantify 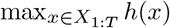 in practice outside of benchmarking exercises.

## Experiments

### Toy data experiment

We begin by demonstrating the overall problem setup on toy data on an input space *x* ∈ [0, 1]. We consider 5 objectives overall – 3 that act as the *useful* objectives with maxima around that of a meta-objective at 1/4 given by sin 2*πx*, max(0, sin 2*πx*), and sin 2*π*(*x* – 0.05) and 2 that disagree and act as the *non-useful* objectives given by 2*x* and – 2*x*. Each objective is augmented with noise (Supplementary Note 2A). Note that on real data we do not know *a priori* which objectives are useful^2^. Further, the meta-objective is not specified – it may be linear, non-linear, and not necessarily a function of the heuristic objectives – it simply needs a maximum at 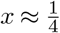.

Samples from each of these functions can be seen in Figure 3A (blue points). The overall Pareto front (orange points) spans almost the entire region including samples at the very right where one of the non-useful linear objective functions has its maximum. However, when applied to this toy problem, MANATEE quickly begins acquiring samples around the joint maxima of the three useful objective functions (red points). Indeed, tracing the inclusion probabilities *p*(λ*_k_* = 1 |**B***_k_*) across the iterations (Figure 3B) demonstrates how MANATEE learns to upweight objectives 1-3 while downweighting 4-5. This demonstrates than when we do not know *a priori* which objectives to trust, we may still recover a region of high utility when the Pareto front spans the full space of conflicting objectives.

**Fig. 3.**
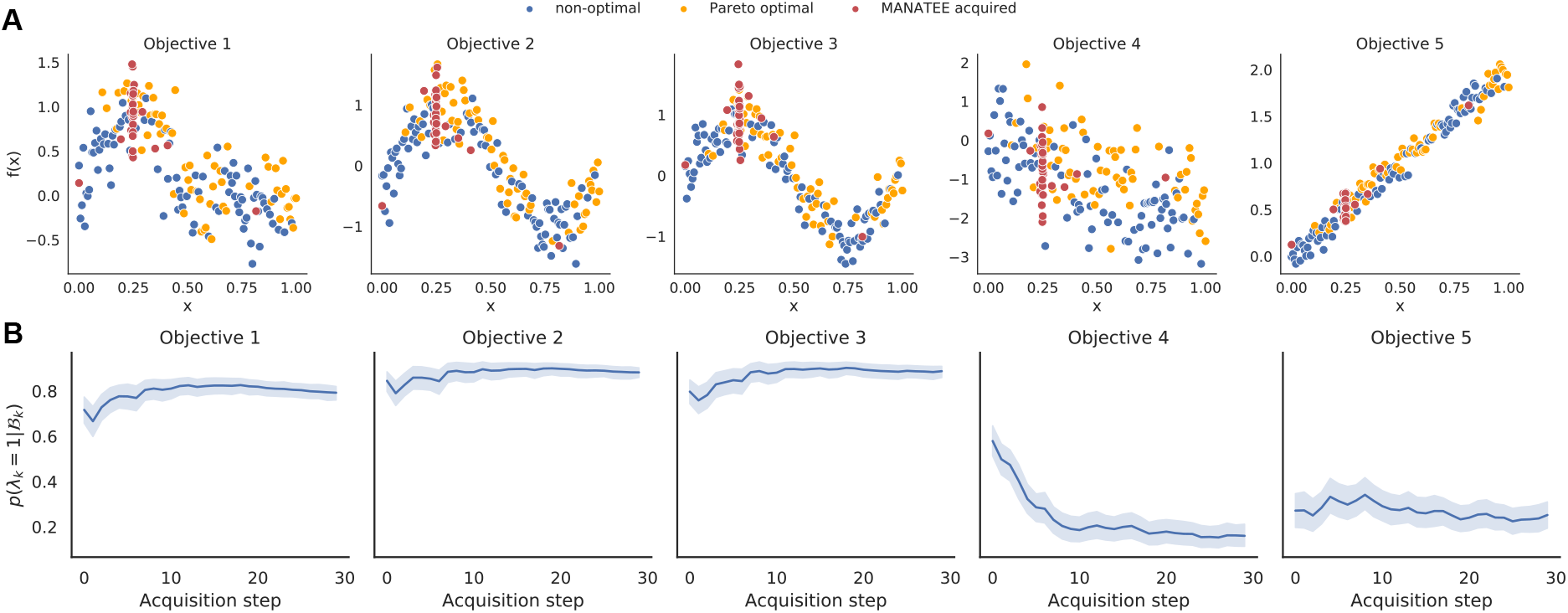
**A** Samples of toy data for the 5 objectives, including those on the Pareto front (orange) and otherwise (blue), along with points acquired by MANATEE-SA (red). **B** Inclusion probabilities for each of the objectives as a function of acquisition step.

### Imaging Mass Cytometry cofactor selection

We next apply MANATEE to the selection of cofactors for Imaging Mass Cytometry (IMC) data, a new technology that can measure the expression of up to 40 proteins at subcellular resolution in tissue sections (25). In the analysis of mass cytometry data, a cofactor *c* is frequently used to normalize the data (26, 27) via the transformation 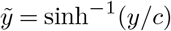. However, to our knowledge no systematic approach exists to set the cofactor and it is typically left as a user-specified parameter.

Here, we consider the standard workflow, where (i) the expression data is normalized with a given cofactor *c* and (ii) the data is clustered using standard methods with the “best” cofactor being the one that leads to the most biologically relevant cellular populations^3^. Given that this problem in general has no notion of “test accuracy” with respect to which we could optimize the cofactor, we instead suggest a number of heuristic objectives based around maximizing the correlation of cluster-specific mean expression of known protein marker combinations. For example, the proteins CD19 and CD20 are highly expressed in B lymphocyte cells and lowly expressed in all others. Therefore, if a clustering correctly separates B cells from others, the correlation between the mean CD19 and CD20 expression in each cluster should be high as the proteins should either be co-expressed or both not expressed (at the origin), as demonstrated in Supplementary Note 6A. We can apply this logic to a range of cell type markers to construct our set of heuristic objectives (Supplementary Note 6B).

To quantify the ability of each clustering to uncover biologically relevant populations, we use expert annotated cell types from (28) and assess cluster overlap with the adjusted Rand index (ARI) and normalized mutual information (NMI), which for this experiment form the overall meta-objectives in line with prior benchmarking efforts of singlecell clustering (29, 30). Note that this is in general unavailable for the analysis of newly generated data and we would *only* have access to the correlation (heuristic) objectives.

The results comparing MANATEE to the alternative methods are shown in Table 1. On the metric of cumulative regret, which as above, is most relevant for the problem setup at hand, MANATEE-SA outperforms the alternative approaches. On full and Bayes regret, MANATEE performs comparably with the baselines. On cumulative regret, *q*NEHVI is comparable to random acquisition, suggesting that consistently acquiring close-to-optimal solutions over > 5 noisy objectives is challenging even for approximate hypervolume computation. Interestingly, we find that random scalarization exhibits strong performance on several measures, which may be understood by the fact that the scalarized objective 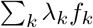 naturally places high weight on regions where many objectives agree, mimicking a similar scenario to our inter-objective agreement criterion.

**Table 1.** Results for IMC cofactor optimization experiment. CR: cumulative regret, FR: full regret; BR: Bayes regret. M-SA: MANATEE with scalarized acquisition, M-AS: MANATEE with acquisition of scalarized function, RA: random acquisition, RS: random scalarization. ARI: adjusted Rand index, NMI: normalized mutual information. Values are mean (s.d.).

| Method | ARI |  |  | NMI |  |  |
| --- | --- | --- | --- | --- | --- | --- |
|  | CR | FR | BR | CR | FR | BR |
| M-SA | **0.018(0.005)** | **0.003(0.003)** | **0.008(0.006)** | **0.019(0.009)** | **0.002(0.005)** | **0.008(0.009)** |
| M-AS | 0.022(0.009) | 0.004(0.010) | 0.009(0.010) | 0.028(0.015) | 0.007(0.016) | 0.014(0.016) |
| RA | 0.045(0.001) | 0.024(0.013) | 0.031(0.008) | 0.065(0.003) | 0.026(0.013) | 0.036(0.009) |
| RS | 0.021(0.004) | **0.003(0.004)** | **0.008(0.005)** | 0.025(0.006) | 0.003(0.006) | **0.008(0.007)** |
| *q*NEHVI | 0.043(0.004) | 0.011(0.007) | 0.021(0.011) | 0.061(0.007) | 0.011(0.007) | 0.026(0.017) |
| *q*NParEGO | 0.037(0.005) | **0.003(0.003)** | 0.018(0.012) | 0.049(0.009) | 0.005(0.003) | 0.023(0.017) |

We further performed ablation experiments of each behaviour and found that no single behaviour drives the performance (Supplementary Note 1D). We also performed crossvalidation on data splits to demonstrate that MANATEE does not overfit to a given dataset (Supplementary Note 1C).

### Single-cell RNA-seq highly variable gene selection

Single-cell RNA-sequencing (scRNA-seq, see (31) for an overview) quantifies whole-transcriptome gene expression at single-cell resolution. A key step in the analysis of the resulting data is selection of a set of highly variable genes (HVGs) for downstream analysis, typically taken as the “top *x*%” (32), but there are no systematic or quantitative recommendations for selecting this proportion (33). Therefore, we apply MANATEE to this problem following a clustering workflow similar to the IMC experiment, but by varying the proportion of HVGs used for the analysis and keeping all other clustering parameters fixed. We again propose a number of co-expression based heuristics (Supplementary Note 6C) and augment these with measures of cluster purity (mean silhouette width, Calinski and Harabasz score, Davies-Bouldin score) previously used in scRNA-seq analysis (1).

For these workflows, no general ground truth clustering or cell types are available. However, a new technology called CITE-seq can simultaneously quantify both the RNA and surface protein expression at single-cell level (34). Given that cell types are traditionally defined by surface protein expression (35), we use a clustering of the surface protein expression alone as the ground truth following existing work (36). The concordance with this clustering acts as the meta-objective in this experiment, which we benchmark the proposed approaches against. We supply each method with the heuristic objectives above and benchmark the gene proportion acquisitions by contrasting the resulting clusterings with the surface protein-derived ground truth using ARI and NMI as metrics. Once again, these represent only two possible choices of meta-objective and there are many more we could design, highlighting the prevalence of heuristic objectives in the field.

The results are shown in Table 2. As above, our main focus is on cumulative regret since when deploying in a real-world scenario, we would not have access to the meta-objective. MANATEE performs favourably on cumulative regret compared to the other approaches, though has higher full and Bayes regret. Overall, this demonstrates our method to be a promising approach to tackle hyperparameter optimization on real, noisy datasets and achieve competitive performance compared to existing baselines and state-of-the-art methods.

**Table 2.** Results for scRNA-seq HVG selection optimization experiment. CR: cumulative regret, FR: full regret; BR: Bayes regret. M-SA: MANATEE with scalarized acquisition, M-AS: MANATEE with acquisition of scalarized function, RA: random acquisition, RS: random scalarization. ARI: adjusted Rand index, NMI: normalized mutual information. Values are mean (s.d.).

| Method | ARI |  |  | NMI |  |  |
| --- | --- | --- | --- | --- | --- | --- |
|  | CR | FR | BR | CR | FR | BR |
| M-SA | 0.119(0.024) | 0.048(0.008) | 0.053(0.008) | **0.114(0.030)** | 0.014(0.014) | 0.022(0.015) |
| M-AS | **0.117(0.027)** | 0.051(0.014) | 0.059(0.014) | 0.117(0.035) | 0.023(0.015) | 0.034(0.019) |
| RA | 0.191(0.022) | **0.045(0.007)** | 0.060(0.014) | 0.201(0.027) | 0.020(0.008) | 0.037(0.017) |
| RS | 0.128(0.017) | **0.045(0.006)** | **0.051(0.007)** | 0.116(0.019) | **0.011(0.003)** | **0.019(0.008)** |
| *q*NEHVI | 0.182(0.054) | 0.046(0.013) | 0.086(0.061) | 0.190(0.067) | 0.021(0.008) | 0.068(0.076) |
| *q*NParEGO | 0.152(0.035) | **0.045(0.006)** | 0.075(0.032) | 0.152(0.040) | 0.013(0.010) | 0.055(0.043) |

## Discussion

A common theme here is the subjectivity of parameter setting in biological data analysis workflows. Setting these often involves no heuristic objectives at all, simply relying on an iterative data exploration to find a parameter combination that “works”. Even when heuristic objectives are involved – such as in the benchmarking analyses of scRNA-seq workflows – the precise choice of which objectives to include is fundamentally subjective too.

It is important to note that our proposed approach does not remove subjectivity from the analysis. Many important steps, including the chosen behaviours 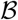 and their conditional inclusion distributions 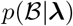 are set by the user. Therefore, it abstracts the subjectivity by a level, changing the question from *“which objectives should I use to benchmark my method?”* to *“what would the behaviour of a good objective function be?”*. Given that no link is assumed between the specified heuristic objectives and the true meta-objective and that the choice of desirable objective behaviours is given as example only, we make no optimality claims about the ability to explore the Pareto front. We note that our approach may be used to optimize parameters in ethically dubious bioinformatics analyses, such as genetic testing of embryos, and strongly caution any such use, emphasizing the need for a thorough ethical review process.

There are several extensions that would serve as future steps. We have only considered 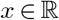, but this could be extended to 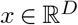 for *D* > 1. Similarly, there is much current research in BO methods over both continuous and categorical domains (37), which may better suit the parameter space of scRNA-seq analysis pipelines (1). Finally, much research in BO centres on the incorporation of user input and expert opinions to guide optimization (38, 39). While we have explicitly considered the opposite problem – where *a priori* it is not known which objectives should be upweighted – there could be situations where both approaches could be integrated. For example, an expert may provide ratings for the results of each scRNA-seq clustering during optimization. In such settings, these ratings could be integrated into our proposed framework by updating the distributions 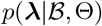 over Θ such that they confer high weights to functions of expert ratings.

## ACKNOWLEDGEMENTS

We gratefully acknowledge funding from the Natural Sciences and Engineering Research Council of Canada (RGPIN-2020-04083) and a CFI/JELF award (950-232944) to KRC, the Canadian Statistical Sciences Institute Ontario Top-up for Postdoctoral Fellows in Data Science to AS, and the Vector Institute Postgraduate Affiliate Award to AS. This research was undertaken, in part, thanks to funding from the Canada Research Chairs Program.

## CODE AVAILABILITY

Code implementing MANATEE used for the analyses can be found in https://github.com/camlab-bioml/2022_manatee_paper_analyses.

## Footnotes

1 Commonly referred to as *tasks*, we here refer to them as *objectives* given the application.

2 Otherwise only useful objectives would be included and standard MOBO procedures applied.

3 All parameters of the clustering procedure are held constant across cofactors to allow for fair comparison.

