## Supplementary Information for "Multi-objective Bayesian Optimization with Heuristic Objectives for Biomedical and Molecular Data Analysis Workflows"

### Supplementary Note 1: Implementation details

**A. Method hyperparameters.** Hyperparameters for all methods are summarized in Table 3.

| Method | Parameter | Value | Explanation |
|---|---|---|---|
| MANATEE | $p(\lambda_k = 1)$ | 0.5 | Prior over binary $\lambda_k$ is set as Bernoulli(0.5) |
| | $\pi_1, \pi_0$ | $0.75, 0.25$ | Bernoulli hyperparameters for $p(B_k^{(3)} \| \lambda_k = i)$ |
| | $\frac{\delta^2 f}{\delta x^2} <$ | -10 | Upper bound to call a max |
| | Min distance | 0.01 | Distance from max to extrema |
| | $l >$ | 0.1 | Kernel lengthscale constraint |
| | $\sigma^2$ | 1 | Kernel variance |
| | $\epsilon^2 >$ | 0.01 | Observation noise variance constraint |
| MANATEE RS | UCB $\beta_t$ | $0.125 \log(2t+1)$ | Set as in (15) |
| | GP fits | 5 | Model inits and fits at each acquisition |
| | GP fit re-tries | 20 | Max re-tries to fit model at each acquisition |
| | Acquisition samples | 100 | Initial samples from acquisition function |
| M-SA | Line search function | `strong_wolfe` | LBFGs optimizer arg |
| M-AS | Schedule | `CosineAnnealingLR` | `lr_scheduler` arg |
| | Steps | 300 | Adam optimizer steps |
| | MC samples | 500 | MC samples to approximate AS |
| | MC samples | 10 | MC samples to evaluate AS for initialization |
| $q$NEHVI $q$NParEGO | `MC_SAMPLES` | 128 | QMC sampler arg, set as in (21) |
| | `max_retries` | 20 | `botorch.fit_gpytorch_model` arg |
| | `batch_range` | $(0, -1)$ | QMC sampler arg, set as in (21) |
| | GP model | `KroneckerMultiTaskGP` | Set as in (21) |
| | `BATCH_SIZE` | 1 | Points to acquire |
| | `NUM_RESTARTS` | 20 | Optimization re-starts, set as in (21) |
| | `RAW_SAMPLES` | 1024 | Acquisition samples, set as in (21) |
| | `batch_limit` | 5 | `optimize_acqf` arg, set as in (21) |
| | `maxiter` | 200 | `optimize_acqf` arg, set as in (21) |
| $q$NEHVI | reference point | sample minimum | Lower HV bound |
| | `alpha` | $10^{-8+\#objectives}$ | approximate partitioning level |

**Table 3.** Method hyperparameters used in the experiments. M-SA: MANATEE with scalarized acquisition, M-AS: MANATEE with acquisition of scalarized function, RS: random scalarization.

**MANATEE** MANATEE (with both AS and SA acquisition functions) was implemented with PyTorch v. 1.9.0 (40) and `gpytorch` v. 1.6.0 (41) for the Gaussian process model and inference. Optimization was performed with the LBFGS optimizer. At every acquisition step, the model was initialized and fit to to the current training set 5 times and the model with the highest log-likelihood was kept. If fitting failed, the process would be re-tried a maximum of 20 times before halting. Optimization of the acquisition function was initialized with the maximum of 100 random samples. Above implementation details also apply to the random scalarization (RS) baseline. For MANATEE, maxima of the posterior mean were identified by computing the first derivative of the posterior mean, finding its zeros, and computing the second derivative at those locations. A candidate was declared a maximum not at boundary if its second derivative was less than -10 and if the candidate was at least 0.01 units away from the range extrema.

**MANATEE-SA** Optimization of the acquisition function was performed with the LBFGS optimizer with line search.

**MANATEE-AS** The second term of the AS acquisition function (see Supplementary Note 3) was approximated with 500 MC samples. Optimization of the acquisition function was performed with the Adam optimizer with cosine annealing. Optimization of the acquisition function was initialized with the maximum of 100 random samples. When evaluating AS on random samples for optimization initialization, 10 MC samples were used for each AS evaluation.

***q*NEHVI** The $q$NEHVI approach was implemented with `botorch` v. 0.6.1.dev37+g4f0a2889 (42). The development version was used to facilitate usage of $q$NEHVI with the `KroneckerMultiTaskGP` model. Implementation closely followed the tutorial on multi-objective Bayesian optimization (21). Batch size was set to 1. `fit_gpytorch_model` was called with `max_retries` set to 20. Other parameters were set following the tutorial. Reference point was set to the minimum of the initial acquired points. To accommodate $> 5$ objectives, we used approximate hypervolume computation by setting the `alpha` parameter according to the heuristic based on the number of objectives as proposed in (20).

***q*NParEGO** The $q$NParEGO approach was implemented with `botorch` v. 0.6.1.dev37+g4f0a2889 (42). Implementation closely followed the tutorial (21). Batch size was set to 1. `fit_gpytorch_model` was called with `max_retries` set to 20. Other parameters were set following the tutorial.

| Experiment | Parameter | Value |
|---|---|---|
| Toy | Number of initial points | 5 |
| | Number of acquisitions | 30 |
| | Replicates | 100 |
| | $x_{min}$ | 0 |
| | $x_{max}$ | 1 |
| | Number of objectives | 5 |
| IMC | Number of initial points | 5 |
| | Number of acquisitions | 35 |
| | Replicates | 98 |
| | $x_{min}$ | 1 |
| | $x_{max}$ | 100 |
| | Number of objectives | 7 |
| scRNA-seq | Number of initial points | 5 |
| | Number of acquisitions | 36 |
| | Replicates | 100 |
| | $x_{min}$ | 0.01 |
| | $x_{max}$ | 0.5 |
| | Number of objectives | 9 |

**Table 4.** Parameters of the experimental procedure.

##### B. Experimental setup.

Parameters for all experimental procedures are summarized in Table 4.

**Toy experiment** The Pareto front on toy objectives was computed with the OApackage (43). The initial dataset contained 5 training points at random locations and MANATEE-SA performed 30 acquisition steps. The experiment was repeated 100 times. The range of the optimized parameter $x$ was between 0 and 1. Toy objectives are described in Supplementary Note 2A.

**IMC experiment** At each acquisition step, data was normalized with the acquired cofactor value and clustered. Mean marker expression in each cluster was computed on the data normalized with cofactor=1. The co-expression objective values were computed as a Pearson correlation between the mean expression of a marker pair across clusters. The experiment was repeated 98 times. The range of the optimized cofactor value was between 1 and 100. The overall meta-objective maximum for ARI and NMI, used to compute regrets, was set as the maximum of ARI/NMI values computed from all acquisitions by all methods (MANATEE-SA, MANATEE-AS, RA and RS baselines, $q$NEHVI, $q$NParEGO), including behaviour ablation methods (MANATEE-SA with leave-one-out behaviour). $q$NEHVI returned an error and completed fewer than the total of 35 acquisitions on 49 runs; $q$NParEGO returned an error and completed fewer than the total of 35 acquisitions on 80 runs, other methods completed all acquisitions on all runs. The objectives are described in Supplementary Note 6B.

**scRNA-seq experiment** At each acquisition step, data was subsetted to the top highly variable genes according to the acquired proportion value and clustered. The co-expression objective values were computed as a Pearson correlation between the mean expression of a marker pair across clusters. The experiment was repeated 100 times. The range of the optimized highly variable gene proportion was between 0.01 and 0.5. Unsupervised cluster purity metrics were computed on the PCA transform of the normalized scRNA-seq data. The overall meta-objective maximum for ARI and NMI, used to compute regrets, was set as the maximum of ARI/NMI values computed from all acquisitions by all methods (MANATEE-SA, MANATEE-AS, RA and RS baselines, $q$NEHVI, $q$NParEGO), including behaviour ablation methods (MANATEE-SA with leave-one-out behaviour). $q$NEHVI returned an error and completed fewer than the total of 36 acquisitions on 63 runs, $q$NParEGO returned an error and completed fewer than the total of 36 acquisitions on 56 runs, other methods completed all acquisitions on all runs. The objectives are described in Supplementary Note 6C.

| Method | IMC | | scRNA-seq | |
|---|---|---|---|---|
| | Runtime | $n$ | Runtime | $n$ |
| M-SA | 576.000(164.662) | 98 | 667.490(219.465) | 100 |
| M-AS | 8222.224(65.969) | 98 | 13895.700(3755.607) | 100 |
| RA | 62.735(0.827) | 98 | 101.540(2.726) | 100 |
| RS | 477.418(13.980) | 98 | 662.190(203.874) | 100 |
| $q$NEHVI | 605.388(396.560) | 49 | 1174.162(515.633) | 37 |
| $q$NParEGO | 1492.056(340.741) | 18 | 2153.114(663.802) | 44 |

**Table 5.** Runtimes of all methods on two experiments. M-SA: MANATEE with scalarized acquisition, M-AS: MANATEE with acquisition of scalarized function, RA: random acquisition, RS: random scalarization. Values are mean (s.d.) in seconds, computed over $n$ complete runs.

Experimental results were tracked with Weights & Biases (44). Reported regrets include acquisitions from incomplete runs for $q$NEHVI and $q$NParEGO. Table 5 shows average runtimes of all methods.

**C. Cross-validation experiments.** In both experiments, subsampled and processed data (as described in Supplementary Note 2) was divided in 70/30% train/test splits in 5-fold cross-validation, where the parameter was optimized with MANATEE-SA only on train and cumulative regrets were computed in train and test. To ensure sufficient data, CITE-seq data was subsampled to 3000 cells in the cross-validation experiments. For the highly variable gene selection experiment, the same set of highly variable genes (computed according to the proportion value acquired on train at each step) was used when computing regrets on test. For both experiments, the initial dataset contained 5 training points at random locations and MANATEE-SA performed 10 acquisitions. Each cross-validation experiment was repeated 100 times. When computing regrets, the overall meta-objective maximum for ARI and NMI was set as the maximum of ARI/NMI values computed from the acquisitions by MANATEE-SA in each run and each fold. For each run, cumulative regret was averaged over folds. Table 6 shows cross-validation cumulative regrets averaged over runs on train and test for both experiments.

| | |
|---|---|
| IMC ARI train | $0.015 \pm 0.003$ |
| IMC ARI test | $0.017 \pm 0.003$ |
| IMC NMI train | $0.021 \pm 0.005$ |
| IMC NMI test | $0.023 \pm 0.005$ |
| scRNA-seq ARI train | $0.093 \pm 0.015$ |
| scRNA-seq ARI test | $0.088 \pm 0.019$ |
| scRNA-seq NMI train | $0.126 \pm 0.024$ |
| scRNA-seq NMI test | $0.128 \pm 0.026$ |

**Table 6.** 5-fold cross-validation mean cumulative regret on train and test data splits. IMC denotes the IMC cofactor optimization experiment, scRNA-seq denotes the highly variable gene proportion optimization experiment. ARI: adjusted Rand index, NMI: normalized mutual information.

**D. Behaviour ablation experiments.** In leave-one-out behaviour experiments, MANATEE-SA was run without one of each behaviours at the time. Table 7 shows results for the IMC cofactor selection experiment, with each row indicating regrets for MANATEE-SA without said behaviour. Table 8 similarly shows results for the scRNA-seq highly variable gene selection experiment.

| Ablated | ARI | | | NMI | | |
|---|---|---|---|---|---|---|
| | CR | FR | BR | CR | FR | BR |
| Explainability | 0.016(0.004) | 0.003(0.003) | 0.007(0.004) | 0.017(0.006) | 0.002(0.005) | 0.007(0.007) |
| Inter-obj agreement | 0.018(0.005) | 0.003(0.005) | 0.008(0.006) | 0.020(0.009) | 0.003(0.008) | 0.008(0.010) |
| Max not at boundary | 0.018(0.007) | 0.004(0.006) | 0.008(0.007) | 0.020(0.012) | 0.004(0.011) | 0.009(0.012) |

**Table 7.** Results for the behaviour ablation experiments for IMC cofactor optimization. CR: cumulative regret, FR: full regret; BR: Bayes regret. ARI: adjusted Rand index, NMI: normalized mutual information. Values are mean (s.d.).

| Ablated | ARI | | | NMI | | |
|---|---|---|---|---|---|---|
| | CR | FR | BR | CR | FR | BR |
| Explainability | 0.114(0.020) | 0.047(0.006) | 0.054(0.008) | 0.106(0.028) | 0.013(0.008) | 0.023(0.013) |
| Inter-obj agreement | 0.118(0.025) | 0.047(0.006) | 0.054(0.009) | 0.110(0.030) | 0.013(0.007) | 0.023(0.013) |
| Max not at boundary | 0.113(0.022) | 0.047(0.010) | 0.054(0.011) | 0.105(0.028) | 0.016(0.011) | 0.025(0.015) |

**Table 8.** Results for the behaviour ablation experiments for scRNA-seq highly variable gene selection. CR: cumulative regret, FR: full regret; BR: Bayes regret. ARI: adjusted Rand index, NMI: normalized mutual information. Values are mean (s.d.).

#### Supplementary Note 2: Data processing

**A. Toy data.** The 5 objectives in the toy experiment had the following functional forms all defined on $x \in [0,1]$:

1. $y_1(x) = \max(0, \sin 2\pi x) + \epsilon$, $\epsilon \sim \mathcal{N}(0, 0.3^2)$
2. $y_2(x) = \sin 2\pi(x - 0.05) + \epsilon$, $\epsilon \sim \mathcal{N}(0, 0.3^2)$
3. $y_3(x) = \sin 2\pi x + \epsilon$, $\epsilon \sim \mathcal{N}(0, 0.3^2)$
4. $y_4(x) = -2x + \epsilon$, $\epsilon \sim \mathcal{N}(0, 0.8^2)$
5. $y_5(x) = 2x + \epsilon$, $\epsilon \sim \mathcal{N}(0, 0.1^2)$

**B. IMC data.** IMC data and expert annotated ground truth clustering used in the experiments come from (28). Data was randomly subsampled to 5000 cells and the heavy metal markers that weren't conjugated to antibodies were removed. Data was clustered with the `scikit-learn` (45) k-means algorithm with $k = 10$.

**C. CITE-seq data.** CITE-seq data used in these experiments come from (34) retrieved using the `SingleCellMultiModal` Bioconductor R package with data version 1.0.0. Cell surface antibody expression was normalized using the `logNormCounts` from the `scuttle` R package (46) and clustered using Seurat v. 4.1.0 (47) with top 10 principal components as input and resolution parameter set to 0.8. Intra-cellular single-cell RNA-seq data was filtered for genes with at least 100 reads and further processed with `scanpy` (48). Data was randomly subsampled to 1000 cells (except for cross-validation experiment, where it was subsampled to 3000 cells) and normalized using `pp.normalize_total` with `target_sum=1e4`, `pp.log1p`, and `pp.scale`. `scanpy` was used to select highly variable genes, compute the neighbourhood graph with 10 neighbours and top 40 principal components, compute the PCA decomposition with default arguments, and compute Leiden clustering with resolution parameter set to 0.8.

#### Supplementary Note 3: Derivations of acquisition functions

**A. Expectation of scalarization of the single-objective acquisition function of objectives (SA).** We define the SA acquisition function as $\mathbb{E}_{p(\boldsymbol{\lambda}|\mathcal{B})}\left[s_{\boldsymbol{\lambda}}(\text{acq}_{\text{UCB}}(\mathbf{f}(x)))\right]$ and derive the following expression:

$$\begin{aligned}
\mathbb{E}_{p(\boldsymbol{\lambda}|\mathcal{B})}\left[s_{\boldsymbol{\lambda}}(\boldsymbol{\mu}(x) + \sqrt{\beta}\boldsymbol{\sigma}(x))\right] &= \mathbb{E}_{p(\boldsymbol{\lambda}|\mathcal{B})}\left[\sum_{k=1}^{K} \lambda_k(\mu_k(x) + \sqrt{\beta}\sigma_k(x))\right] \\
&= \sum_{k=1}^{K} \mathbb{E}_{p(\boldsymbol{\lambda}|\mathcal{B})}\left[\lambda_k\right](\mu_k(x) + \sqrt{\beta}\sigma_k(x)) \\
&= \sum_{k=1}^{K} p(\lambda_k = 1|\mathbf{B}_k)(\mu_k(x) + \sqrt{\beta}\sigma_k(x))
\end{aligned}$$

**B. Expectation of single-objective acquisition function of the scalarized objectives (AS).** We define the AS acquisition function as $\mathbb{E}_{p(\boldsymbol{\lambda}|\mathcal{B})}\left[\text{acq}_{\text{UCB}}(s_{\boldsymbol{\lambda}}(\mathbf{f}(x)))\right]$ and wish to derive the following expression:

$$\mathbb{E}_{p(\boldsymbol{\lambda}|\mathcal{B})}\left[\text{acq}_{\text{UCB}}\left(\sum_{k=1}^{K}\lambda_k f_k(x)\right)\right]$$

First, notice that,

$$(\lambda_1 f_1(x),...\lambda_K f_K(x)) \sim \mathcal{N}\left(\boldsymbol{\lambda}\boldsymbol{\mu}(x), \boldsymbol{\lambda}^T\boldsymbol{\lambda}\Sigma(x)\right),$$

where $\boldsymbol{\mu}(x)$ is the posterior mean and $\Sigma(x)$ is the posterior covariance of $\mathbf{f}$ evaluated at some $x$.
Then, their sum is distributed as:

$$\sum_k \lambda_k f_k(x) \sim \mathcal{N}\left(\sum_k \lambda_k \mu_k(x), \sum_k \lambda_k^2 \Sigma_{kk}(x) + 2\sum_{1\leq i<j\leq K}\lambda_i\lambda_j\Sigma_{ij}(x)\right)$$

Now, we use this to derive:

$$\begin{aligned}
&\mathbb{E}_{p(\boldsymbol{\lambda}|\mathcal{B})}\left[\text{acq}_{\text{UCB}}\left(\sum_{k=1}^{K}\lambda_k f_k(x)\right)\right]\\
&= \mathbb{E}_{p(\boldsymbol{\lambda}|\mathcal{B})}\left[\sum_k \lambda_k\mu_k(x) + \sqrt{\beta}\cdot\sqrt{\sum_k \lambda_k^2\Sigma_{kk}(x) + 2\sum_{1\leq i<j\leq K}\lambda_i\lambda_j\Sigma_{ij}(x)}\right]\\
&= \sum_k p(\lambda_k = 1|\mathbf{B}_k)\mu_k(x) + \sqrt{\beta}\cdot\mathbb{E}_{p(\boldsymbol{\lambda}|\mathcal{B})}\left[\sqrt{\sum_k \lambda_k^2\Sigma_{kk}(x) + 2\sum_{1\leq i<j\leq K}\lambda_i\lambda_j\Sigma_{ij}(x)}\right]
\end{aligned}$$

We approximate the expectation term with $S$ samples of $\lambda_k^s \sim p(\lambda_k|\mathbf{B}_k)\ \forall\ k$ and $s = 1,\ldots,S$:

$$\begin{aligned}
&\mathbb{E}_{p(\boldsymbol{\lambda}|\mathcal{B})}\left[\sqrt{\sum_k \lambda_k^2\Sigma_{kk}(x) + 2\sum_{1\leq i<j\leq K}\lambda_i\lambda_j\Sigma_{ij}(x)}\right]\\
&\approx \frac{1}{S}\sum_{s=1}^{S}\sqrt{\sum_k \left(\lambda_k^s\right)^2\Sigma_{kk}(x) + 2\sum_{1\leq i<j\leq K}\lambda_i^s\lambda_j^s\Sigma_{ij}(x)}
\end{aligned}$$

#### Supplementary Note 4: Derivatives of a multi-output Gaussian process

The posterior mean of a Gaussian process with noisy observations $\mathbf{y}$ at a new location $x_*$ is given by (13):

$$\bar{f}_* = \mathbf{K}(x_*, X)\left(\mathbf{K}(X,X) + \sigma_\epsilon^2 I_N\right)^{-1}\mathbf{y}.$$

For the exponentiated quadratic kernel function, the first derivative of the posterior mean (which is also the mean of the distribution over derivatives of the GP posterior functions) is (49):

$$\begin{aligned}
\frac{\delta \bar{f}_*}{\delta x_*} &= \frac{\delta \mathbf{K}(x_*, X)}{\delta x_*}\boldsymbol{\alpha} = -\Lambda^{-1}\tilde{X}^T\left(\mathbf{K}(x_*, X)^T \odot \boldsymbol{\alpha}\right)\\
\boldsymbol{\alpha} &= \left(\mathbf{K}(X,X) + \sigma_\epsilon^2 I_N\right)^{-1}\mathbf{y}\\
\tilde{X} &= [x_* - x_1, \ldots, x_* - x_N]^T.
\end{aligned}$$

Above, $\tilde{X}$ is an $N \times D$ matrix for $D$-dimensional input $x$ and $\odot$ represents element-wise multiplication.
For $D = 1$ case, let us label the lengthscale kernel hyperparameter as $l$ and derive the second derivative of the posterior mean to be:

$$\frac{\delta^2 \bar{f}_*}{\delta x_*^2} = \left(-\frac{1}{l^2}\mathbf{K}(x_*, X) + \frac{1}{l^4}\tilde{X}^T \odot \tilde{X}^T \odot \mathbf{K}(x_*, X)\right)\boldsymbol{\alpha}$$

For a multi-output Gaussian process with $K$ objectives, the first and second derivatives of the posterior mean are given by the above Equations with arranging observations $\mathbf{y} = [y_{11}, \dots, y_{N1} \dots y_{1K}, \dots, y_{NK}]^T$ and $\tilde{X} = [x_* - x_1, \dots, x_* - x_N \dots x_* - x_1, \dots, x_* - x_N]^T$ as $KN \times 1$ vectors.
The multi-output kernel is defined as $\mathbf{K}^{\mathrm{IO}} \otimes \mathbf{K}$ and the additive noise term is $D \otimes I_N$, where $D$ is the diagonal matrix with task-specific observation noises. The resulting derivatives are $K$-dimensional, with each element representing the task-specific derivative w.r.t. $x_*$.

#### Supplementary Note 5: Proofs of theorems

*Theorem 1*: If $\mathbb{E}_{p(\lambda_k|\mathbf{B}_k)}[\lambda_k] > 0 \,\forall k$, the solution to $\max_x \mathbb{E}_{p(\boldsymbol{\lambda}|\mathcal{B})} s_{\boldsymbol{\lambda}}(\mathbf{f}(x))$ lies on the Pareto front of $\mathbf{f}$.

*Proof:* Let $\psi_k := \mathbb{E}_{p(\lambda_k|\mathbf{B}_k)}[\lambda_k] > 0$. Then by linearity of expectation $\mathbb{E}_{p(\boldsymbol{\lambda}|\mathcal{B})} s_{\boldsymbol{\lambda}}(\mathbf{f}(x)) = \sum_k \mathbb{E}_{p(\lambda_k|\mathbf{B}_k)}[\lambda_k] f_k(x) = \sum_k \psi_k f_k(x)$ is monotonically increasing in all $f_k$ and the result follows from (15). ■

*Theorem 2*: For some $p(\boldsymbol{\lambda}|\mathcal{B})$, any point $x^*$ on the Pareto front of $\mathbf{f}$ is reachable as a maximizer of $\mathbb{E}_{p(\boldsymbol{\lambda}|\mathcal{B})} s_{\boldsymbol{\lambda}}(\mathbf{f}(x))$.

*Proof:* Let $x^*$ be the maximizer of $\sum_k \alpha_k f_k(x)$ with $\alpha_k > 0 \,\forall k$ which is by definition a point on the Pareto front of $\mathbf{f}$. Then $x^*$ is also the maximizer of $\sum_k \delta\alpha_k f_k(x)$ for a constant $\delta > 0$. We may set $\delta$ such that $\delta < \frac{1}{\max_k \alpha_k}$ and so $0 < \delta\alpha_k < 1 \,\forall k$ and thus $\delta\alpha_k$ may be expressed as the expectation of a Bernoulli R.V. $\lambda_k$ for some $p(\lambda_k)$. ■

#### Supplementary Note 6: Cluster mean co-expression as a heuristic

**A. Overview.** The cluster mean co-expression heuristic is demonstrated in Figure 4. After clustering the single-cell data, we can consider the expression of two proteins which should be *markers* for a given cell type, i.e. they should either both be co-expressed or not expressed. An example of this is shown in Figure 4 (right). Consequently, the correlation in the cluster means is high. Conversely, if the clustering does not capture the cell types well, the correlation in the cluster means will be low (Figure 4 left). The opposite logic applies if two proteins should be mutually exclusively expressed: the correlation of cluster mean expression should be minimized.

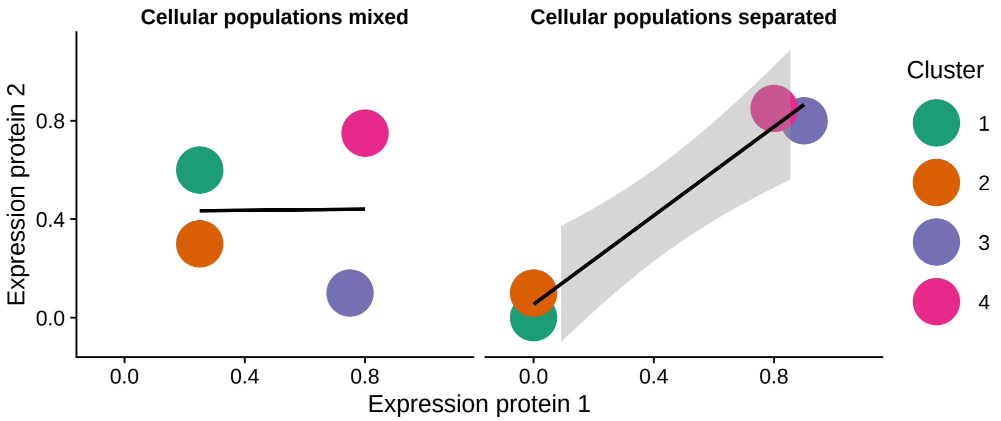


**Fig. 4.** Mean expression of two example proteins after two sets of clustering. Left: the cluster means poorly separate into double-positive and double-negative populations as would be expected if the two proteins are markers for the same cell type. Left: the ideal situation, where clusters only co-express both proteins simultaneously or not at all.

**B. IMC experiment.** The protein pairs used to construct co-expression heuristic objectives are listed in Table 9.

| Protein pair | Co-expression direction | Cell type |
|---|---|---|
| E-Cadherin, pan-Cytokeratin | + | Epithelial |
| CD45, CD20 | + | B cell |
| CD45, CD68 | + | Myeloid |
| CD45, CD3 | + | T cell |
| Vimentin, Fibronectin | + | Stromal cell |
| CD19, CD20 | + | B cell |
| pan-Cytokeratin, Cytokeratin 5 | + | Basal epithelial |

**Table 9.** Co-expression of marker pairs used as objectives in the IMC cofactor selection experiment.

| Gene pair | Co-expression direction | Cell type(s) |
|---|---|---|
| *CD3E, CD4* | + | Regulatory T cell |
| *CD3D, CD8A* | + | Cytotoxic T cell |
| *PTPRC, CD68* | + | Myeloid |
| *CD19, MS4A1* | + | B cell |
| *CD3D, CD68* | - | T/myeloid |
| *CD68, MS4A1* | - | Myeloid/B |

**Table 10.** Co-expression of marker pairs used as objectives in the scRNA-seq highly variable gene selection experiment.

**C. scRNA-seq experiment.** The gene pairs used to construct co-expression heuristic objectives are listed in Table 10.
